# Temporal Dynamics of Activity in Default Mode Network Suggest a Role in Top-Down Processing for Trial Responses

**DOI:** 10.1101/2023.01.08.523152

**Authors:** D. Mastrovito, C. Hanson, S. Hanson

**Affiliations:** Allen Institute for Brain Science, Seattle WA; Department of Psychology Rutgers University, Newark NJ

## Abstract

The default mode network (DMN) is a collection of brain regions including midline frontal and parietal structures, medial and lateral temporal lobes, and lateral parietal cortex. Although there is evidence that the network can be subdivided into at least two subcomponents, the network reliably exhibits highly correlated activity both at rest and during task performance. Current understanding regarding the function of the DMN rests on a large body of research indicating that activity in the network decreases during task epochs of experimental paradigms relative to inter-trial intervals. A seeming contradiction arises when the experimental paradigm includes tasks involving autobiographical memory, thinking about one’s self, planning for the future, or social cognition. In such cases, the DMN’s activity increases and is correlated with attentional networks. Some have therefore concluded that the DMN supports advanced human cognitive abilities such as interoceptive processing and theory of mind. This conclusion may be called into question by evidence of correlated activity in homologous brain regions in other, even non-primate, species. Thus, there are contradictory findings related to the function of the DMN that have been difficult to integrate into a coherent theory regarding its function. Using data from the Human Connectome Project, we explore the temporal dynamics of activity in different regions of the DMN in relation to stimulus presentation. We show that generally the dorsal portion of the network exhibits only a transient initial decrease in activity at the start of trials that increases over trial duration. The ventral component often has more similarity in its time course to that of task-activated areas. We propose that task-associated ramping dynamics in the network are incompatible with a task-negative view of the DMN and propose the dorsal and ventral sub-components of network may rather work together to support bottom-up salience detection and subsequent top-down voluntary action. In this context, we re-interpret the body of anatomical and neurophysiological experimental evidence, arguing that this interpretation can accommodate the seeming contradictions regarding DMN function in the extant literature.

## I. Introduction

The default mode network (DMN) is a collection of brain regions including midline frontal and parietal structures, medial and lateral temporal lobes, and lateral parietal cortex (Buckner, Andrews-Hanna, Schacter, 2008; Mazoyer et al., 2001; Raichle et al., 2001; Shulman et al., 1997). The Default Mode Network (DMN) reliably exhibits a decrease in blood-oxygen-level-dependent (BOLD) signal intensity, across a wide variety of task conditions (Shulman et al., 1997; Mazoyer et al., 2001; Laird et al., 2009), commonly referred to as deactivation. Generally, during task performance, DMN activity is negatively correlated with activity in regions of the executive control network (Greicius et al., 2003; Fair et al., 2007, Seeley et al., 2007) and dorsal attention networks (Corbetta & Shulman, 2002, Fox et al., 2006), regions that increase in activity during task performance. This set of observations led to the description of the DMN as a “task-negative” network (Fox et al. 2005). However, significant correlation within the network persists even during task-associated deactivation (Greicius et al., 2003; Greicius & Menon, 2004). In addition, not all experimental paradigms evoke decreases in DMN regions (Spreng et al. 2010). Increases, rather than decreases, in DMN activity are evoked for tasks involving autobiographical memory (Cabeza, 2004), self-referential processing (Gusnard et al., 2001; Buckner & Carroll, 2007; Andrews-Hanna et al., 2010b), theory of mind (Amodio & Frith, 2006; Carrington & Bailey, 2009), semantic processing (Binder, 2009), task-switching (Crittenden et al., 2015), and planning for the future (Baker et al., 1996; Spreng et al., 2010). This set of observations suggests an alternative hypothesis that the DMN supports cognitive abilities such as interoceptive processing and theory of mind, abilities typically ascribed specifically to humans. However, in spite of the extensive evolutionary expansion of association cortex, which makes up much of the DMN, a well-organized intrinsically coherent network resembling the human DMN has been identified in other species including non-human primates Mantini et al., 2011), mice (Stafford et al., 2014), cats (Popa et al., 2009) and even rats (Lu et al., 2007; Upadhyay et al., 2011). It is therefore clear that none of our current hypotheses about the function of this network are able to account for all of the observations in the extant literature. That is, the DMN cannot both be task-negative and be associated with performance of a specific subset of tasks. In addition, the meaning and function of correlated, albeit lower intensity, activity during tasks that evoke deactivation in the network are not addressed by either of the existing theories of DMN function.

The function of human brain networks are often inferred from their increases in activity during task-based fMRI paradigms. Typically, analysis of task-based fMRI data involves averaging over blocks of similar trials. This approach, however, obscures information about the temporal dynamics of activity in each region. Evidence from both task and resting-state fMRI using various analytical techniques indicates that there may be subcomponents of the DMN with differing temporal dynamics (Laird et al. 2009; Andrews-Hanna et al. 2010; Zuo et al. 2010; Smith et al. 2012; Hsu et al. 2016; Uddin et al., 2009), though the differential function of these subcomponents is not well understood. Beyond mean signal increases during task performance, insights into the function of these subcomponents may be gained from closer attention to the regions whose activity is most often associated with them. For example, one region that is reliably identified as part of the DMN is Brodmann area 32 in the anterior cingulate (Shulman et al., 1997) (Figure 4). Although it is often overlooked, considerable evidence from both animal and human literature indicates that this region is reliably predictive of action initiation (Mukamel, et al. 2011; Srinivasan et al. 2013) and with action selection including error monitoring (Luu et al., 2000), conflict resolution (Carter et al., 1998). These observations call into question the notion that the network as a whole does not participate in cognitive tasks, even when, on average, relative decreases in network activity are observed. Indeed, it has also been shown that BOLD signal measured from the DMN during task performance contains task-specific patterns that can be used to distinguish task conditions (Crittenden et al., 2015). However, the dynamics of activity in subcomponents of the DMN in relationship to task performance remains relatively unexplored. The following work aims to address this. Borrowing from the neurophysiological toolbox, we characterize the peristimulus activity in different regions of the DMN during task epochs of the Human Connectome Project (HCP) visual processing, and social cognition paradigms. Our results indicate that each of the DMN regions show task-specific fluctuations, albeit with different temporal dynamics in different parts of the network: with ventral DMN preferentially active during the beginning of trials and dorsal portions of the network showing an initial decrease at the beginning of trials followed by a steady increase over trial duration. In light of these observations, we reconsider the existing literature on DMN function, and propose that the ventral and dorsal subcomponents of the DMN may work together to orchestrate bottom-up and top-down volitional control of cognitive processes.

**Figure 1.**
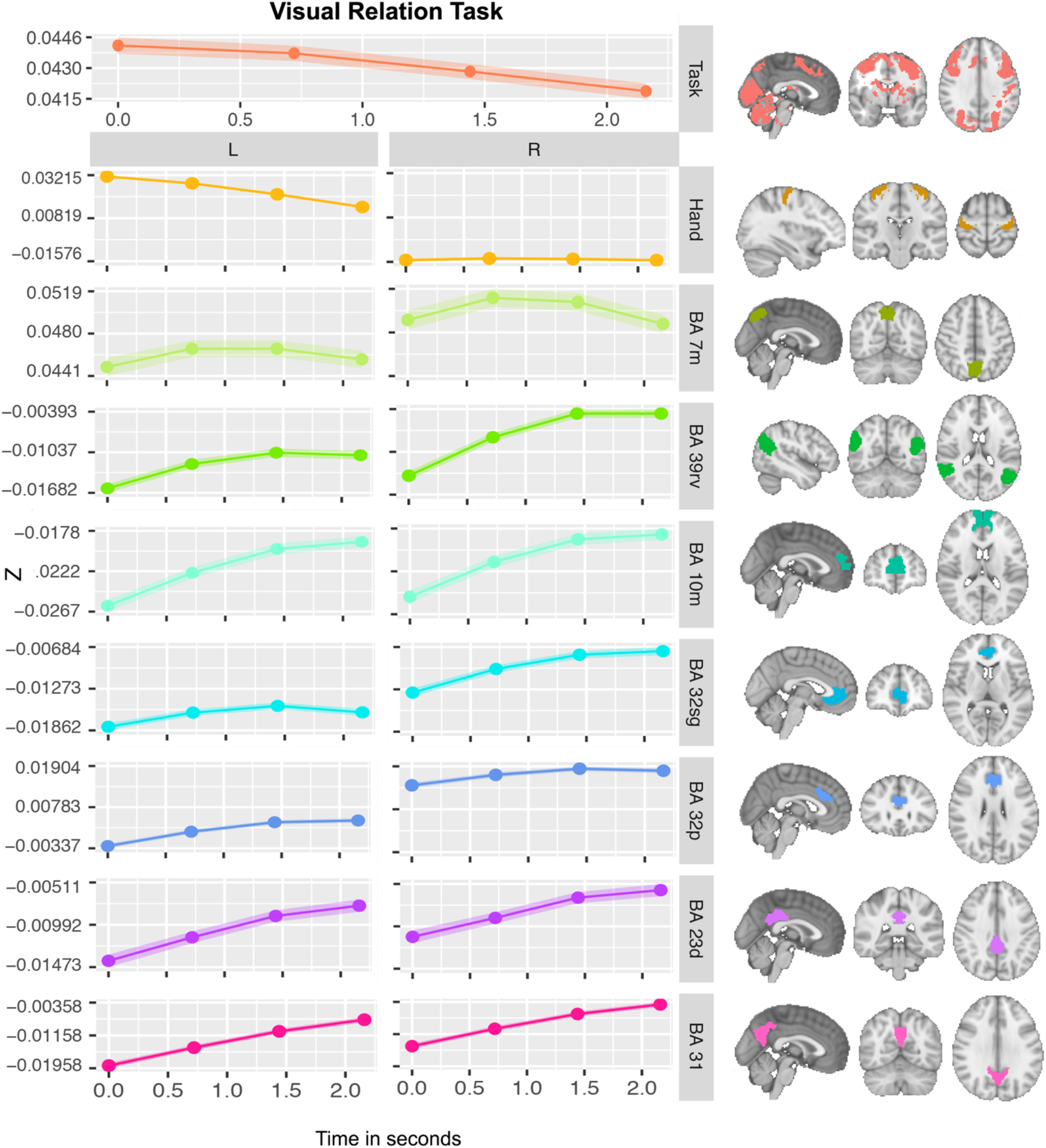
Relational-Visual Processing DMN Peristimulus Activity. Average peristimulus activity over all conditions of the visual processing paradigm. Intensity values are centered on each regions mean value during rest. Thus, negative values represent an intensity below that of inter-trial intervals and positive values above. Shaded region indicates standard error of the mean. L Hand corresponds with the left motor cortical area associated with movement of the right hand. All regions of the network exhibit a decrease in activity relative to inter-trial intervals at the beginning of the trial, but also exhibit increasing signal over the course of the trial.

**Figure 2.**
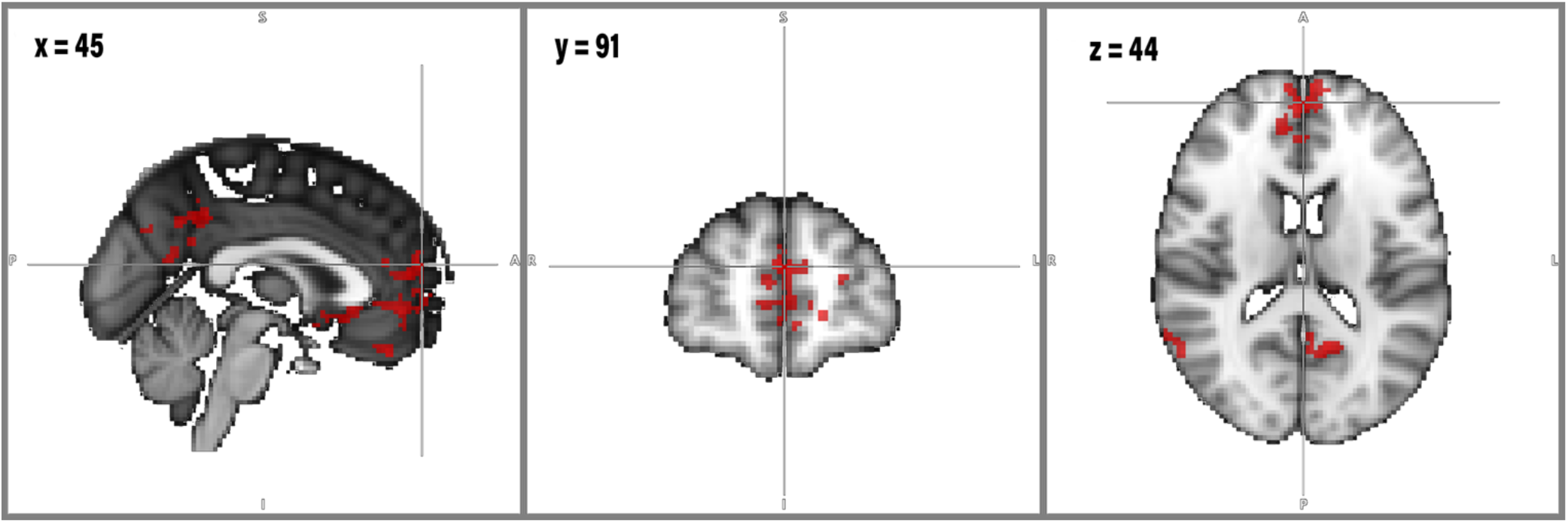
DMN exhibits ramping of activity during relational processing paradigm. Group average of contrast of ramping model and square model of the experimental design.

**Figure 3.**
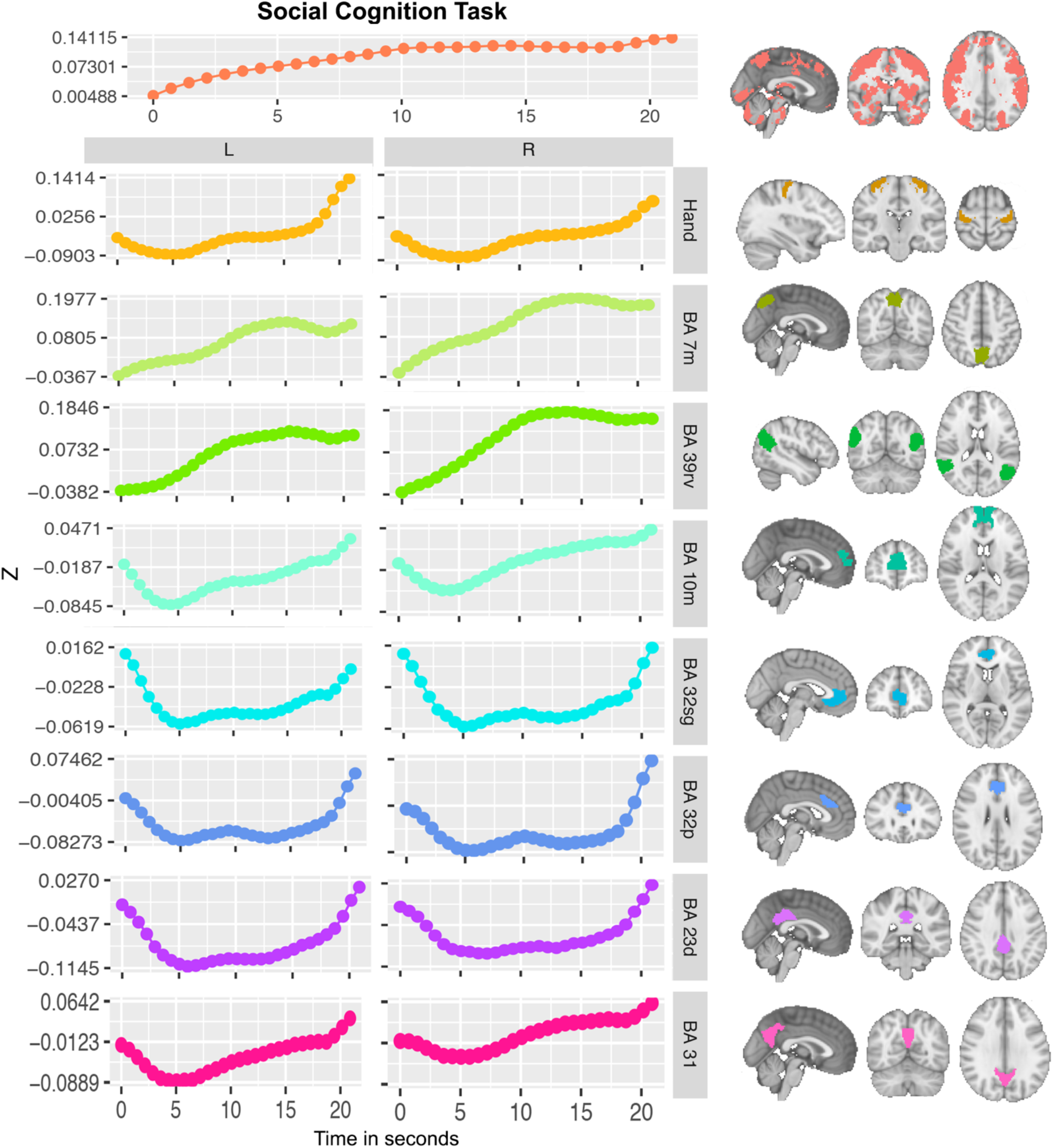
Social Processing DMN Peristimulus Activity. Average peristimulus activity for the social condition only of the social cognition paradigm. The network appears to have 3 distinct patterns of activity, 1 in the ventral component, and the others in the dorsal component, one in the anterior cingulate, where ramping activity is not observed over the course of the trial, but increases coincide with the timing of trial responses, while all other regions of the dorsal network show a decrease in signal at the beginning of the trial followed by a steady increase over the duration of the trial.

**Figure 4.**
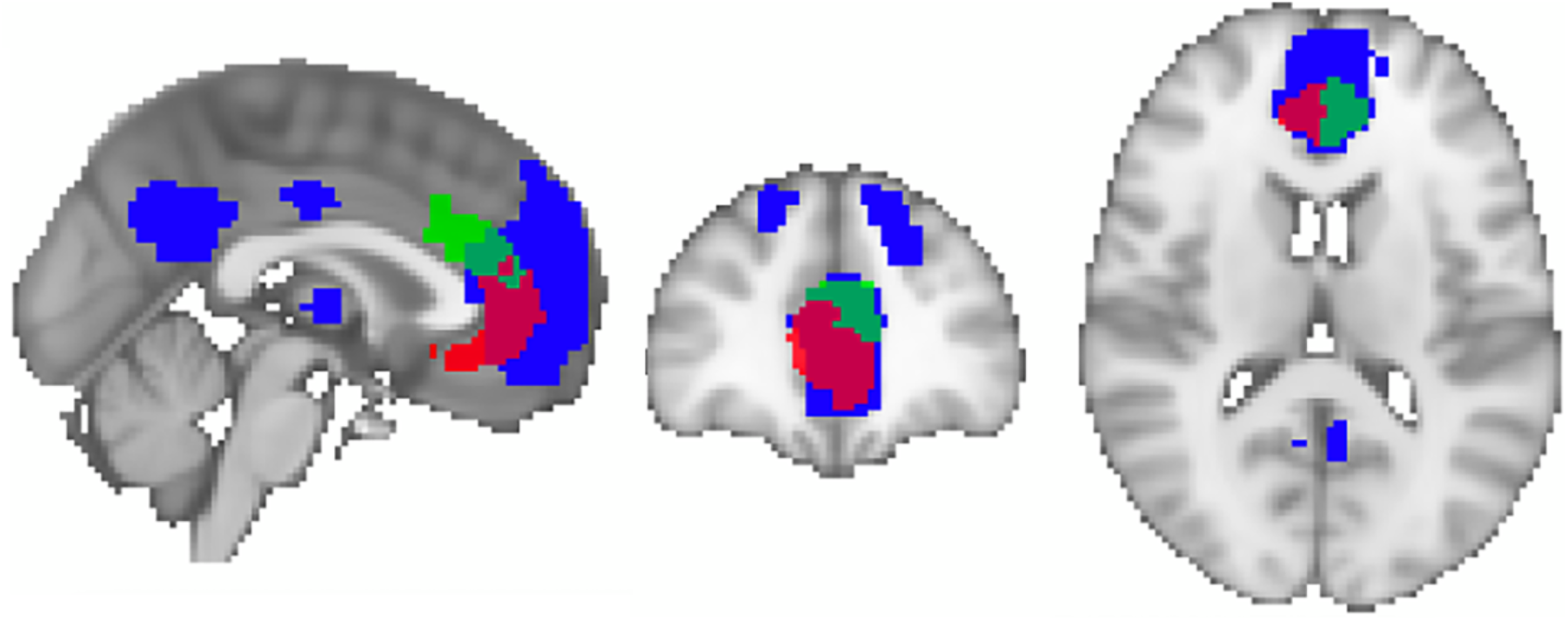
Overlap of Dorsal Default Mode Network and Brodmann Area 32. Dorsal DMN shown in blue from the Willard functional atlas (Richiardi & Altmann, 2015) defined by multimodal independent component analysis combining resting state fMRI and post-mortem gene expression. BA 32 pregenual (green) and BA 32 subgenual (red) defined from Brainnetome atlas (Fan et al., 2016)

## II. Methods

### Data

Minimally preprocessed functional MRI was acquired from the Human Connectome Project (HCP) healthy young adult data set (Glasser et al., 2013). A subset of the available data used for this work was selected based on the criteria that subjects completed both the Social and Relational MRI protocols. This subset included 1004 subjects. Details of the HCP minimal preprocessing pipeline have previously been described (Glasser et al. 2013), but consist of motion correction using six degrees of freedom (DOF) linear image registration with FSL’s FLIRT, registration to a single band anatomical reference image and corrections for B0 field inhomogeneities using FSL’s topup (Andersson et al., 2003). Transformations for registration and distortion correction (gradient nonlinearity distortion, motion correction and EPI distortion) are concatenated and applied in a single transform with spline interpolation. Data is then masked and normalized to a 4D whole brain mean of 10,000. The resulting images are in 2mm Montreal Neurological Institute (MNI) space. 91 left-handed subjects were later excluded from analysis in order not to confound results based on differences in lateralization associated with handedness. The remaining sample included 913 subjects: 499 females, and 414 males; aged between 22 and 36 years of age: mean +- SD = 28.75 +- 3.72 years. The following post-hoc analyses were carried out on the resulting sample, which included two fMRI runs per subject, per task-paradigm.

The HCP fMRI protocol includes seven task paradigms (van Essen et al. 2013). Two of these were chosen for this analysis for 1) their use of an experimental block design and 2) their purported differential engagement of the DMN (exhibiting task-evoked increases vs decreases). These were the social cognition, and the relational processing respectively. The social cognition paradigm is a theory of mind task. Theory of mind is a cognitive ability that is believed to engage the DMN and to evoke increases in DMN activity (Fletcher et al., 1995). The relational task involves visual information processing. Visual processing tasks were among the first shown to evoke decreases in DMN activity (Shulman, 1997). Therefore, we aimed to see how the temporal dynamics of activity in the DMN differed across these two tasks. The details of each of these tasks have previously been described (Barch et al., 2013a) but a basic overview is provided here. The social cognition task is based on Frith– Happe animations of social and random interactions developed for the purpose of studying theory of mind (Castelli et al., 2000; Wheatley et al., 2007). In 20 second video clips, geometric figures (squares, circles, and triangles) move either randomly on the screen or in a manner suggesting social interaction. After each video, participants judged whether the objects interacted with each other or moved randomly. Each of two task runs had five video blocks (two interaction blocks and three random blocks in one run, three interaction and two random in the other run) and five fixation blocks (15 seconds each).

The relational task is a visual information processing paradigm involving stimuli with different shapes and textures (Smith et al., 2007). In each trial, participants are presented with two pairs of objects, one at the top of the screen and one at the bottom of the screen. In the relational condition, subjects first decide whether the pair at the top of the screen differs in shape or texture and then decide whether the bottom pair differs along the same dimension (shape or texture) as the top pair. In a matching condition, subjects are presented with one pair at the top of the screen and a single image at the bottom of the screen. A cue with the word “shape” or “texture” in the middle of the screen instructs subjects to decide if the image at the bottom of the screen matches either image at the top of the screen on that dimension. In the relational condition, the stimuli are presented for 3500 ms with a 500ms ITI, with four trials per block. In the matching condition, the stimuli are presented for 2800 ms, with a 400 ms ITI and five trials per block. In each of two runs there are three 18 second relational blocks, three 18 second matching blocks and three 16 second fixation blocks.

### Definition of Regions of Interest

#### DMN Regions

We chose to use the Brainnetome atlas (Fan et al., 2016), because its parcellation is based on Brodmann areas, which allowed us to use regional definitions that are more easily related with those used in the animal literature. Regions of interest (ROIs) for the DMN were identified within the Brainnetome atlas with the use of a separately generated and previously published functional atlas consisting of 14 functional networks (Shirer et al. 2012). Brainnetome regions overlapping with the ventral DMN (vDMN) and dorsal DMN (dDMN) as defined in the functional atlas included bilateral Brainnetome regions: dDMN: BA 10m (medial), BA 32p (pregenual), BA 31; vDMN: BA 7m (medial), BA 39rv (rostroventral), and BA 23d (dorsal). These ROIs were used for all analysis. Mean time series for each of these DMN ROIs were extracted, and detrended to remove linear trends in the time series for each subject and task.

#### Task-activated Regions

For each task, a GLM analysis was conducted using FLS’s Feat (Smith et al. 2004; Jenkinson et al. 2012) to identify brain regions exhibiting task-evoked activations on average over task conditions at the group level. First, voxel-wise multiple regression of experimental block designs was performed at the single subject level (Woolrich et al. 2001). This analysis included pre-whitening, spatial smoothing using a Gaussian kernel of 4 mm full width half max (FWHM) and high pass temporal filtering with a cutoff of 200 seconds. Activation map Z statistic images were created for single subjects with an initial cluster threshold of Z > 3.1 and corrected cluster significance with threshold of p < 0.05 (Worsley, 2003). For each paradigm, a second level GLM was fitted using a fixed effects model combining the two runs for each subject. This resulted in an average activation map for each subject and condition of each experimental paradigm. For each paradigm, group level analysis was then carried out using a mixed-effects model (FLAME1) (Beckmann et al., 2003; Woolrich et al., 2004). The resulting group-level Z statistic maps with corrected cluster significance threshold of p < 0.05, were indicative of voxels that on average exhibit increases in BOLD signal associated with experimental task epochs. For each task, these voxels were then used to define a single “Task” ROI that served to represent the time course of task-activated brain activity. Mean time-series for this “Task” ROI were extracted from the GLM derived voxels and detrended for each task and subject.

#### Bilateral Hand-associated Motor ROI

In order visualize DMN activity relative to the timing of subject responses, an ROI was created for voxels associated with button-press responses of the subjects’ right hand at the end of each trial. This ROI was created using data from the social cognition paradigm, because its experimental design included response epochs that could be modeled as a separate condition. As we only include right-handed subjects, a group-level activation map for the response condition was used to identify voxels in the left-lateralized motor cortex associated with the button press response, using FLS’s Feat as described above. Mean time series of these voxels was extracted, and detrended to remove linear trends in the time series for each subject. For comparison, time series from the same voxels in the contralateral motor cortex were also extracted.

### Peristimulus Activity

In neurophysiological studies, peristimulus histograms are often used to visualize changes in neuronal firing rate in relation to an external event or stimulus. Analogously in fMRI, peristimulus plots can be used to visualize task-evoked changes in BOLD signal to qualitatively characterize their dynamics. In order to understand the dynamics of signal change in each of the DMN regions over experimental epochs, visualizations of peristimulus activity in each of the DMN regions were created. Time points associated with individual trials within experimental epochs were identified using the available E-Prime logs, and modeled with a gamma hemodynamic response function (HRF). For each subject and ROI, average signal during rest epochs was used as a baseline. ROI time series were centered on this value and normalized by their standard deviation over the entire scan so that positive values indicate signal intensity greater than that during inter-trial intervals. The resulting average time series in each ROI was then extracted and averaged across subjects. To account for different experimental block lengths without discarding data, each trial was linearly interpolated to match the number of time points to the mean trial length over all trials for all subjects. Peristimulus plots for each paradigm were created by averaging data over trials for all subjects including two runs each (1826 scans in total for each task): relational task = 45,588 trials; mental condition of the social cognition paradigm = 4565 trials. For each task, peristimulus activity associated with task performance was visualized using a single ROI which included 1000 voxels sampled from the group-level activation map for that task. To further establish the significance of the signal change over individual trials, paired t-tests (one-tailed) were subsequently performed comparing the final time points of each trial across subjects to the first time points (Relational task: Table 2). For the social paradigm, this comparison was between the time point of mean minimum intensity early in the trial to the last time point before trial response period: Table 3 (all p-values were Bonferroni corrected for multiple comparisons).

**Table 1.** Average BOLD Percent Change over Experimental Blocks

|  | Percent Change |  |  |  |
| --- | --- | --- | --- | --- |
| Task | DMN |  | Task activated regions |  |
|  | Mean | Max | Mean | Max |
| Relational | -0.46% +- 0.20 | 3.0% +- 1.47 | 0.73% +- 0.19 | 17.26% +- 4.73 |
| Social | -0.37% +- 0.19 | 1.37% +- 0.74 | 0.65% +- 0.23 | 16.75% +- 4.99 |

**Table 2.** Relational Cognition BOLD Percent Change in DMN regions over Trials

| dDMN | Percent Change % | t-statistic | p value |
| --- | --- | --- | --- |
| **Anterior** |  |  |  |
| BA 10m L | 27.04 | 15.30 | 9.16 e-52 |
| BA 10m R | 26.03 | 13.80 | 2.7 e-42 |
| BA 32p L | 252.55 | 12.59 | 2.56 e-35 |
| BA 32p R | 27.05 | 6.24 | 3.74 e-9 |
| BA 32sg L | 11.30 | 4.39 | 8.40 e-24 |
| BA 32sg R | 43.77 | 10.27 | 2.69 e -13 |
| **Posterior** |  |  |  |
| BA 23d L | 44.71 | 9.34 | 8.46 e -24 |
| BA 23d R | 47.74 | 7.59 | 2.69 e -13 |
| BA 31 L | 59.05 | 18.78 | 1.77 e -77 |
| BA 31 R | 71.13 | 18.04 | 1.56 e-71 |
| **vDMN** |  |  |  |
| BA 7m L | 1.57 | 1.11 | 2.26 |
| BA 7m R | -0.72 | -0.52 | 11.9 |
| BA 39rv L | 32.70 | 9.32 | 1.04 e-19 |
| BA 39rv R | 67.06 | 16.86 | 1.26 e-62 |
| **Task** | -5.12 | -7.25 | 17 |
| **Left Motor Cortex** | -54.03 | -26.27 | 17 |
| **Right Motor Cortex** | -0.38 | -0.07 | 8.98 |

**Table 3.** Social Cognition BOLD Percent Change in DMN regions over Trials

| **dDMN** | **Percent Change %** | **t-statistic** | **P value** |
| --- | --- | --- | --- |
| **Anterior** |  |  |  |
| BA 10m L | 311.44 | 10.06 | 9.6 e-13 |
| BA 10m R | 658.87 | 7.72 | 1.1 e-13 |
| BA 32p L | 17949.42 | 3.72 | 0.0017 |
| BA 32p R | 858.12 | 0.24 | 6.9 |
| BA 32sg L | -154.42 | 5.47 | 4.0 e-7 |
| BA 32sg R | 67.69 | 2.84 | 0.039 |
| **Posterior** |  |  |  |
| BA 23d L | 226.41 | 5.99 | 1.8 e-08 |
| BA 23d R | 293.38 | 2.30 | 0.19 |
| BA 31 L | 274.70 | 8.75 | 2.1 e-17 |
| BA 31 R | 821.16 | 5.51 | 3.1 e-7 |
| **vDMN** |  |  |  |
| BA 7m L | 461.32 | 6.24 | 3.8 e-9 |
| BA 7m R | 795.84 | 5.49 | 3.6 e-7 |
| BA 39rv L | 413.74 | 7.58 | 3.2 e-13 |
| BA 39rv R | 598.86 | 0.16 | 7.4 |
| **Task** | 2113.64 | 0.15 | 7.5 |
| **Left Motor Cortex** | 475.02 | 4.04 | 0.00046 |
| **Right Motor Cortex** | 374.96 | 3.24 | 0.01 |

### GLM Modeling of Task-evoked Ramping Activity

The peristimulus activity of DMN regions during the relational task, as described above, suggested that activity within DMN regions might exhibit near-linear increases over trial durations. To determine if this activity was statistically significant over trials at the group level, a custom design file was created with a linear ramping function over trials. That is, instead of the typical square wave or box-car design with ones during experimental epochs and zeros representing inter-trial intervals, a triangle wave design was used. As we were specifically interested in DMN activity, which we expect to be maximal during inter-trial epochs, inter-trial rest periods were labeled with ones. The first TR associated with each trial was marked by a zero and subsequently linearly ramped to a value of one over the trial duration. The end of each trial was determined by utilizing the subject response times for each trial. To determine if the ramp model was a better fit for DMN activity than a square wave with ones during inter-trial intervals and zeros during the task, a contrast between the two designs was performed in a first level fixed-effects analysis using FSL Feat for each of 100 subjects randomly selected from the larger sample. A higher-level fixed-effects analyses was conducted to estimate the average across runs within participants. Finally, a mixed-effects analysis, treating subjects as random effects, was carried out using FSL FLAME1. A cluster-based thresholding with a Z threshold of 3.1 and p value 0.5 was used at each level.

## III. Results

### Peristimulus Activity in the DMN Activity is Region Specific and Does Not Exhibit Sustained Suppression

Task-evoked activations are characterized by sustained increase in BOLD signal over blocks of repeated experimental trials. In both experimental paradigms, the corresponding decreases in BOLD signal intensity over experimental blocks in the DMN are relatively small (Table 1). We expected decreases in DMN activity during the relational task paradigm and increases in the social cognition paradigm. However, the overall pattern of task-associated increases and decreases across the network were similar for both experimental paradigms. Visualizing the temporal dynamics of task-evoked signal changes in the DMN revealed that “deactivations” in the DMN are not equal and opposite that of task-activated regions. Rather than a sustained suppression of activity over trials, we found that when a region exhibited task-evoked decreases, these decreases were transient, occurring only at the beginning of experimental trials with steadily increasing intensity over trial durations.

### Relational Paradigm – Visual Processing

Because data from each region was centered on their mean activity during inter-trial intervals, all negative peristimulus intensity values are indicative of activity below that during inter-trial intervals and positive values indicate activity above that of inter-trial intervals. The Relational task paradigm is known to evoke decreases in DMN activity. As expected, we found that, with the exception of BA 7m and BA 32p in right-lateralized anterior cingulate, all other regions of the DMN exhibited some BOLD signal attenuation relative to that during inter-trial intervals (Figure 1). However, it is readily ascertained from the peristimulus activity, where time =0.0 represents the start of the trial, that nearly all regions of the DMN exhibit an initial decrease in signal intensity, below that of resting levels, at the beginning of the trial. However, the BOLD signal in each region subsequently steadily increases over the duration of the trial. Importantly all regions of the network approached baseline levels of activity over the duration of the trial. In this task, BA 7m in the vDMN, seems to differ from the rest of the network, in that the signal there never dropped below baseline levels and rather exhibited sustained increases in activity over the entirety of the trial, similar to that of task-activated brain areas. Calculating the percent change in activity from the beginning of the trial to the end, revealed that regions of the dDMN generally exhibit large increases in activity over the course of individual trials (Table 2), while the unique task-evoked time courses suggest different functional roles for different subcomponents of the network. All regions of the DMN, except BA 7m, were significantly higher at the final time points of each trial than at the beginning of the trial, a pattern that is not seen in regions defined as “task-activated” and identified by square-wave model of the task.

In subsequent GLM analysis, which explicitly modeled ramping activity over individual trials across individuals contrasted against a box-car model, the same regions were identified, as there were significant clusters in all regions of the DMN except bilateral BA 7m and right lateralized BA 32p (Figure 2). The percent overlap (number of voxels) of each cluster with each region of the DMN is listed in Table 3, indicating the largest proportion of voxels better explained by a ramping model in bilateral BA 31 in the posterior cingulate as well as large proportions in the anterior cingulate and BA 10m in medial prefrontal cortex.

### Social Cognition Paradigm

In the social paradigm, ramping activity levels can also be seen in many DMN regions during trials. The ramping behavior is most prominently seen in BA 10m and BA31, while other regions in the dorsal part of the network exhibit a marked increase in activity coincident with the beginning of trial responses (the last few points beyond 20 seconds of video presentation) and with BOLD activity in the motor cortex during response epochs (Figure 3). BAs 32sg, 32p and 23d all show increases in responses in particular at the end of the video presentation at the start of the trial response period. The rest of the dorsal part of the network shows transient decreases early during the trial and subsequent ramping activity for the duration of the trial often exceeding mean intensity during rest periods. The vDMN also exhibits increases in signal intensity over the trial duration, exceeding mean resting levels but these increases seem to plateau toward the end of the trial. The peristimulus activity in BA 7m shows sustained increases in activity in both the social and relational paradigms. Peri-stimulus activity suggests that task-evoked activity in medial parietal cortex has a distinct pattern from that in the cingulate and medial prefrontal cortex. Both experimental paradigms (social and relational) evoked a more sustained increase in parietal cortex. In the social paradigm however, the anterior DMN in the cingulate exhibited trial-associated fluctuations resembling those associated with trial responses and increases in signal strength over the course of trial durations were generally more significant in left-lateralized cortex than right (Table 3). The patterns BAs 10m and 31 resemble that of the relational paradigm with transient decreases at the beginning of trials followed by steady increases over the trial duration.

It is worth noting that in addition to our expectation of differential engagement of the DMN across these tasks: DMN engagement in the social cognition task, and lack of engagement in visual processing task, the two cognitive tasks analyzed here also differ significantly in the length of time associated with a trial. In the social cognition task, a trial lasts more than 20 seconds, and is an order of magnitude shorter in the visual processing task. This difference might account in part for the differences observed in the time courses of the dorsal DMN, particularly in the anterior cingulate. Despite these differences, there nevertheless remains remarkable similarity in the dynamics of activity across these regions in relation to task performance, in that in both paradigms activity in the network gradually increases over the course of the trial. In previous work, similarities in DMN activity (mean decreases across trial blocks) across a wide range of task domains were noted. Here we demonstrate that the network may also exhibit similar temporal dynamics across task domains, dynamics which suggest a greater role in task performance than has been previously appreciated.

## Discussion

In this work, we examined the temporal dynamics of peristimulus activity in each region of the DMN during two experimental paradigms involving visual and social processing. We found that relative decreases in DMN activity do not represent sustained suppression of activity over experimental trials, but rather are indicative of transient decreases in activity specifically at the beginning of each trial within an experimental block. Peristimulus fluctuations in BOLD signal intensity across the DMN are consistent with the timing of individual trials within blocks, in correspondence with the specific experimental design. This corroborates other studies which have previously reported that activity in the DMN contained task-specific information (Vatansever et al., 2015) and raises the intriguing possibility that in fact the DMN is not inactive during task performance as has long been suggested. We further identified subcomponents of the DMN, with distinct temporal dynamics during task performance, suggesting distinct functional roles. The observation of distinct temporal dynamics within dorsal and ventral subcomponents of the network is also consistent with previous studies, particularly of restingstate fMRI, in which similar subcomponents of the DMN with independent time courses were identified (Smith et al. 2012; Bado et al. 2014). The measured peristimulus activity we report supports this subdivision and further suggests specific functions for the two subcomponents of the network. Consistent with work indicating that portions of the medial parietal cortex become uncorrelated with the DMN and correlated with the TPN during task execution (Leech et al., 2011), we found that the ventral subcomponent increased in activity during visual processing and activity in BA 7m in fact did not drop below baseline levels of activity measured during inter-trial intervals, suggesting the involvement of the ventral component of the network during visual processing. Regional decreases in BOLD signal intensity occurred specifically in the dorsal part of the network. These decreases in activity are small relative to task-evoked increases and, more importantly, are transient, occurring specifically at the beginning of individual trials and exhibiting a ramping increase in activity over trial durations. Neurophysiological studies have demonstrated transient changes in neural activity in response to external task demands corresponding to sequential bottom-up and top-down flow of information (Siegel et al., 2015). Furthermore, ramping activity is believed to be a cortical mechanism underlying temporal control of action (Narayanan 2017). Therefore, time courses in the dorsal portion of the network, in contrast, suggest a role specifically in facilitating trial responses. To our knowledge, however, this is the first time ramping activity in the dorsal DMN has been reported and especially during experimental epochs of tasks believed to evoke decreases in DMN activity. The observation calls into serious question the conclusion that the DMN is task-negative, as such a conclusion would require sustained decreases in network activity during task performance that resumes only after task completion. The ramping of activity in the dorsal portion of the network, in combination with sustained increases in the ventral portion of the network, suggests that the DMN may actively support evidence accumulation from task-positive regions in preparation for trial responses. Therefore, we propose an alternative hypothesis regarding the function of the DMN, whereby the two subcomponents of the network work together to determine task-relevant information from externally received, in this case, specifically visually presented information, while the other integrates information from higher order cognitive processing in regions generally thought to be task-positive, in preparation for top-down generated responses. When performing tasks that require processing of external information, the ventral portion of the network increases in activity early. In contrast, the dorsal portion of the network supports self-directed top-down processing, increasing in activity later and ramping over the task duration. In tasks that do not require processing external information - tasks such as thinking about oneself, or planning actions in the future, the DMN exhibits a sustained increase in activity. In this view, the network is not “task-negative”, and the dorsal portion of the network gradually increases in activity during bottom-up processing in preparation for initiating top-down responses. This proposed function of the network provides a parsimonious explanation for DMN function, in that it resolves the apparent contradiction between the network as task-negative on the one hand, and as supporting certain types of cognition on the other. It also provides a functional explanation for previous work indicating that the DMN and task-positive regions can exhibit periods of coupled activity at rest and during task performance (Spreng et al., 2010; Dixon et al., 2017; Dixon et al., 2018). Furthermore, it provides a simple explanation for the prominence of DMN-like activity in infants and other species including rodents, which while they may not engage in self-reflection, are capable of self-directed cognition.

This proposed function of the DMN is consistent with the dual mechanism of cognitive control theory, which posits that there are two main modes of conscious brain activity. The first is a bottom-up sensing mode where the brain acquires information from the environment and processes that information using domain-specific brain areas in combination with brain regions supporting working memory and the required attentional resources. A second mode is engaged when enough information has been acquired that a response can be made, a goal-directed top-down mode. fMRI experiments are typically designed with the goal of identifying regions supporting domain specific processing involved in the cognitive processes of interest. Such studies therefore are primarily measuring the first mode, the externally directed portion of cognitive processing. The two modes likely correspond to the differing roles of the DMN and the TPN, with largely anticorrelated activity in the two as the brain switches between bottom-up and top-down processing. However, the key insight is that the DMN does participate in tasks requiring externally focused attention. First, the vDMN helps to identify salient stimuli, perhaps signaling directly or through subcortical structures, to TP regions that focus should be directed towards salient sensory information; and subsequently the dDMN receives information coming from task-related brain areas, gradually building enough evidence to make a decision. The time course of this activity is transient relative to activity in task-related brain areas, more typically the intended targets of study. In the following sections we present arguments in favor of this view based on anatomical and methodological considerations as well as findings from the animal literature.

### fMRI Experimental Design and Analysis Techniques Obfuscate DMN Function

The standard approach to fMRI analysis, in which average activity during different experimental epochs is subtractively contrasted, obscures the temporal dynamics of the BOLD signal. Average contrastive analysis has helped identify regional functional associations as well as networks of regions that together support specific kinds of cognition. Indeed, the Human Connectome data used in this study, when analyzed in this common manner, identifies decreases in activity in DMN regions (Barch et al. 2013) in the relational processing task. The DMN reliably exhibits, on average, a relative decrease in activity while performing a variety of cognitive tasks (Shulman et al. 1997; Fox et al. 2005), rebounding at the end of experimental trials, leading to the proposal of the DMN as a task-negative network. This is true in event-related (Shannon et al., 2006) as well as block design studies, where it is assumed that post-trial increases in DMN activity support the resumption of internally directed thoughts associated with mind-wandering in the resting state. While it can be onerous to examine the temporal dynamics of activity in all regions of the brain simultaneously, critically, it is precisely these dynamics that distinguish areas of the dorsal DMN from task-negative to exhibiting only transient decreases in activity at the beginning of trials and provides a hint into the function of this poorly understood area of the brain. Further evidence for the view of the task-negative DMN, came from studies suggesting that decreases in DMN activity are required to maintain attention on externally directed tasks and that harder tasks require greater decreases in DMN activity (McKiernan et al. 2001). Many studies report a correlation between task-evoked suppression of DMN activity and task accuracy (Weissman et al., 2006; Kelly et al., 2008; Esterman et al., 2013). Therefore, it is thought that a reduction in DMN activity is required to successfully redirect attention away from internal rumination towards the external environment (Buckner et al., 2008). Others have suggested that decreased activity in the DMN may act to reduce brain activity in regions supporting task-irrelevant functions (Anticevic et al., 2013). While not mutually exclusive with our proposed DMN function, harder tasks typically require more time to process, a critical potential confound that is not typically controlled for. If cumulative preparatory activity in the dorsal DMN were spread out the over this longer period of processing time, it would on average appear to be a greater decrease in signal. Paradoxically, the network is also associated with increased activity during specific cognitive activities such as thinking about oneself, planning for the future and remembering the past. Some have posited that the network may therefore be preferentially involved in internally directed, rather than externally directed, attention. However, there is another important difference between the types of tasks known to evoke increases rather than decreases in the DMN. Tasks such as planning for the future, or self reflection are self-directed tasks that do not require bottom-up processing of external (visual) stimuli. In fact, outside of studies designed to probe self-reflective or social cognition, MRI research is almost always conducted by visually presenting stimuli to subjects and requiring them to subsequently make responses via button box. Therefore, such studies almost universally require a period of bottom-up processing of stimuli prior to top-down driven response. If the dorsal DMN supports accumulation of evidence and top-down driven responses, rather than internally directed attention per se, the most commonly used experimental design in combination with mean subtractive analysis would certainly obfuscate this possibility. In this view, resumption of base-line levels of DMN activity at the end of experimental trials reflect the resumption of self-directed rather than necessarily self-reflective cognitive activities.

### Anatomical Considerations

The DMN is easily identified in resting-state fMRI using ICA or PCA, consisting of midline structures in frontal, orbital and parietal cortices, including regions in medial orbital prefrontal cortex. Perhaps because it is so easily recognizable, beyond identifying the network as a whole, researchers seldom identify the specific regions that make up the network. When one examines the composite regions more closely, one region of the brain that is consistently included in the network is the anterior cingulate, BA32 (Figure 4). The precise function of each of the architectonic subdivisions of BA32 are not yet agreed upon (Stevens 2011 for review). Generally, the subgenual anterior cingulate is believed to be connected to pain-related thalamic nuclei as well as the amgydala (Vogt 2006; Ghashghaei et al 2007). Therefore, the inclusion of the subgenual anterior cingulate can be understood in terms of the DMN’s purported role in rumination and social/emotional processing. In contrast, the pregenual anterior cingulate is more often associated with various aspects of action selection including error monitoring (Luu et al., 2000), conflict resolution (Carter et al., 1998), and action initiation (Srinivasan et al. 2013). It can be seen in Figure 4 that the DMN overlaps substantially with both subdivisions of the anterior cingulate. However, to our knowledge, little work has been done to reconcile the large body of literature on BA32’s role in action initiation and its temporal correlation with activity in the DMN. A large body of neurophysiological literature supports the role of the anterior cingulate in voluntary movement including one study that found voluntary movements can be predicted by changes in neural firing rate in the anterior cingulate (Fried et al., 2011). In fact, there is substantial evidence implicating regions of the DMN in the representation of intention and in initiation of self-generated motor activity. This evidence is derived from lesions in animals and in patients (see Haggard 2008 for review). Most intriguingly, one study of voluntary action in humans found that the content of a subject’s choice could be predicted seconds before the readiness signal in supplementary motor areas, which signal the onset of movement, in regions of the DMN, especially BA 10 (Soon et al. 2008).

The DMN consists of regions that have undergone significant expansion in humans, particularly those in medial prefrontal cortex (BAs 10 and 32) (Ongür & Price, 2000) for which there are no clear homologues in non-human species and whose functional role is therefore less well understood. However, in spite of the extensive evolutionary expansion of association cortex in humans, a well-organized intrinsically coherent network resembling the human DMN has been identified in other species. This observation calls into question the meta-cognitive functions ascribed to the DMN for which there is little evidence other species engage. Studies of resting state fMRI have revealed coherent activity in homologous network regions in mice (Stafford et al., 2014), rats (Lu et al., 2007; Upadhyay et al., 2011) and cats (Popa et al., 2009). Task- associated deactivations in macaques are also similar to those identified in human studies (Mantini et al., 2011). The proposed alternative functional role of the DMN also provides an explanation for the existence of DMN activity in other species in that each engage in willful, self-directed action. Although there is likely no homologue for medial prefrontal cortex and in particular BA 10 of the human brain, as ramping activity has frequently been reported as a strong indication of motor initiation medial frontal structures of the non-human primate and rodent anterior cingulate.

## Conclusion

Combining evidence from the set of tasks that evoke increases in the DMN and observations of the ventral portion of the network’s likely role in perception of salient visual stimuli, the most parsimonious of the currently proposed functions of the DMN is that of a sensory-visceromotor link (Raichle, 2015) that pairs experience with appropriate behavioral and emotional responses (Ongür & Price, 2000). This conceptualization of DMN function is also consistent with the idea that, behaviorally, the DMN may underlie stimulus-independent thought, as suggested by McGuire et al. 1996. Our key insight rests on the observation that the dorsal DMN gradually increases in activity during trials with only a transient decrease at the beginning of each trial. This observation in combination with the above proposed function of the DMN allows for a parsimonious explanation of a variety of seemingly contradictory findings regarding DMN activity, whereby tasks that result in decreases in DMN activity reflect only transient decreases as the subject accumulates relevant task information before making a response and DMN activity in non-human species can be understood regardless of the species’s relative ability to engage in self-reflective meta-cognition. Future work should include replication of these results in experiments designed to explicitly test for ramping activity in DMN regions during task performance. Additionally, in order to more fully understand the dynamics of activity in subcomponents of the DMN, it may be of particular interest in future experiments to explicitly model trial periods in the fMRI experimental design as a linear ramp in order to identify regions that exhibit growing preparatory activity. Such a design, when applied across cognitive domains, would allow us to determine whether dDMN regions exhibit this task-evoked temporal profile in a non-task-specific manner and to establish whether the ramping pattern of activity we report is universally present in tasks known to evoke decreases in DMN activity. Finally, although the time course of dDMN activity suggests a possible role for the dDMN in facilitating task responses, we do not mean to imply that the dDMN directly controls motor activity. It is important to note that there is a rich and extensive literature on ramping activity in the brain (Narayanan 2017) where it is has been suggested that such activity may encode the accumulation of information over time underlying human decision making (Soon et al. 2008). The purpose of this ramping activity in the dDMN during task performance should be a focus of future research.

**Table 4.** Group Level Mean General Linear Model Result. Regions identified with greater parameter estimates in ramp over box-car model. Percent indicates percent of voxel overlap with its associated ROI in the Brainnetome atlas.

| **DMN regions** | **Percent** |  | **Other regions 5 1%** | **Percent** |
| --- | --- | --- | --- | --- |
| BA31_R | 23.82 |  | aSTS_R | 23.24 |
| BA31_L | 19.01 |  | dmPOS_L | 17.87 |
| BA10m_R | 17.32 |  | BA14m_R | 16.98 |
| BA32sg_L | 16.19 |  | vld/vlg_R | 13.94 |
| BA32sg_R | 15.69 |  | BA22r_L | 12.42 |
| BA23d_R | 12.83 |  | BA14m_L | 11.97 |
| BA10m_L | 10.88 |  | BA22r_R | 9.81 |
| BA23d_L | 10.25 |  | dmPOS_R | 9.13 |
| BA39rv_L | 9.26 |  | rpSTS_R | 8.83 |
| BA32p_L | 3.79 |  | vld/vlg_L | 8.81 |
| BA39rv_R | 2.48 |  | BA10l_L | 6.93 |
|  |  |  | cpSTS_R | 6.55 |
|  |  |  | BA4ul_R | 6.23 |
|  |  |  | BA24rv_R | 5.92 |
|  |  |  | aSTS_L | 5.42 |
|  |  |  | BA1/2/3ulhf_R | 5.05 |
|  |  |  | rpSTS_L | 5.02 |

